# N, N-Dimethyltryptamine and harmine formulation shifts metastable topography sequences in the cortex

**DOI:** 10.64898/2025.12.04.692302

**Authors:** Maria Niedernhuber, Dila Suay, Michael Mueller, Milan Scheidegger, Dario Dornbierer, Helena Aicher

**Author notes:** Correspondence concerning this article should be addressed to Niedernhuber, Maria, University of Zurich Department of Psychology Binzmuehlestrasse 14, Box 1 8050 Zurich Switzerland. Author note The authors made the following contributions. Niedernhuber, Maria: Conceptualization, Data analysis, Writing - Original Draft Preparation, Writing - Review & Editing; Suay, Dila: Writing - Review & Editing, Supervision; Mueller, Michael: Writing - Review & Editing, Supervision; Scheidegger, Milan: Writing - Review & Editing, Supervision; Dornbierer, Dario: Writing - Review & Editing, Supervision; Aicher, Helena: Writing - Review & Editing, Supervision, Data Collection.

## Abstract

Classic serotonergic psychedelics are potent modulators of conscious awareness, yet the principles governing their effects on the temporal dynamics of brain activity remain unclear. Dominant theories propose that psychedelics increase the signal diversity and metastability of neural states, but whether this reflects a simple randomization of activity or a more structured reorganization is unknown. Here, we used high-density EEG (n=25) in a double-blind, placebo-controlled study to delineate the effects of an ayahuasca-inspired formulation (intranasal N,N-DMT and buccal harmine) on the syntax of whole-brain functional states. Using microstate analysis, we show that psychedelic states induced by DMT and harmine are characterized by a profound acceleration of neural dynamics, evidenced by a reduced microstate duration and an increased frequency of state transitions. Despite that, the sequence of microstates became less random, as indicated by a higher first-order Markov statistic. This syntactic restructuring involved a systematic de-prioritization of transitions into one state (M2) and a corresponding increase in the prevalence and accessibility of two other states (M3 and M5). These findings reveal that the psychedelic state leads to a syntactically reconfigured state of heightened metastability. By showing that psychedelics both lead to an accelerated exploration of an expanded repertoire of neural states governed by increased syntactic structure, our findings improve our understanding of neural metastability in psychedelic states.

## Introduction

Recent years have witnessed a profound resurgence of interest in using psychedelic compounds to probe the neural mechanisms of consciousness (Rankaduwa & Owen, 2023; Timmermann et al., 2023; Yaden et al., 2021). Psychedelics are psychoactive drugs which produce temporary changes in emotion, perception and cognition not readily explained by altered sensory input or diminished wakefulness (Vollenweider & Kometer, 2010). At a neurobiological level, classic psychedelics bind to serotonin (5-HT2A) receptors which are densely concentrated in frontoparietal brain regions thought to be important for conscious awareness (Beliveau et al., 2017; Dehaene & Changeux, 2011; Sanchez, Hartmann, Fuscà, Demarchi, & Weisz, 2020). There is increasing evidence that psychedelics promote neuroplasticity via 5-HT2A receptor agonism (Ly et al., 2018; Nichols, 2016). As a result, psychedelics are currently being explored for the treatment of neurological and psychiatric conditions (Chi & Gold, 2020; Johnson, Hendricks, Barrett, & Griffiths, 2019; Vollenweider & Preller, 2020).

Classic psychedelics include hallucinogens such as N,N-Dimethyltryptamine (DMT), lysergic acid diethylamide (LSD), mescaline, and psilocybin (Johnson et al., 2019). DMT is a tryptamine which produces a range of altered conscious experiences such as changes in self perception, vivid mental imagery, and even spiritual experiences. Unlike other psychedelic compounds, DMT can lead to a rapid and complete immersion into a separate reality often described as a “breakthrough” (Lawrence, Carhart-Harris, Griffiths, & Timmermann, 2022; Timmermann et al., 2019). Ayahuasca combines the potent psychedelic DMT with a monoamine oxidase inhibitor (MAO-I) to produce a prolonged altered state of consciousness (Domínguez-Clavé et al., 2016; Singleton et al., 2022). Ingestion of DMT with the MAO-I harmine prevents the psychedelic from decomposing in the digestive track. This leads to a psychedelic experience lasting several hours, and involving deep introspection, emotional processing, and visionary states integrated with personal memory (Carbonaro & Gatch, 2016).

According to a recent proposal, psychedelics alter consciousness by shifting the brain into a dynamically flexible, informationally diverse state optimised for global integration (Girn et al., 2023). Based on this view, psychedelic states are enabled by an increase in neural metastability, i.e., the brain’s capacity for fluid transitions between distinct functional states (Tognoli & Kelso, 2014). By making the brain more dynamically flexible, psychedelics might open up access to a much larger repertoire of potential brain states (Girn et al., 2023). Indeed, there are several lines of evidence for this model of increased metastability for psychedelic action (Girn et al., 2023). Key findings include greater variance in network synchrony and more diverse functional connectivity patterns with psilocybin (R. Carhart-Harris et al., 2014; Lord et al., 2019; Tagliazucchi, Carhart-Harris, Leech, Nutt, & Chialvo, 2014), as well as an expanded repertoire of harmonic brain states and a lower energetic barrier for state transitions under LSD (Atasoy et al., 2017; Singleton et al., 2022).

Moment-to-moment neural activity in the resting conscious brain can be described as a sequence of transitions between distinct metastable topographical scalp maps, commonly referred to as EEG microstates (Bréchet & Michel, 2022; Michel & Koenig, 2018). Importantly, these metastable sequences of sensor-space configurations can be clustered into a number of dominant cortical topographies (Michel & Koenig, 2018). Labelled “atoms of thought” (Lehmann, 1990) or “basic building blocks of mental processes”(Lehmann & Michel, 2011), microstates have been suggested as a measure to access fluctuations in the stream of consciousness when the brain is idle (Lehmann, 1990). Evidence shows that the stability of these states and the flexibility of the transitions between them are intimately linked to conscious experience and can be tracked on sub-second timescales (Bréchet & Michel, 2022). Accordingly, microstate dynamics shift in various altered states of consciousness such as meditation (Faber, Travis, Milz, & Parim, 2017; Zanesco et al., 2021), dreaming (Bréchet, Brunet, Perogamvros, Tononi, & Michel, 2020; Diezig, Denzer, Achermann, Mast, & Koenig, 2022), anaesthesia (Hermann et al., 2024; Lapointe, Li, Hudetz, & Vlisides, 2023; Li, Vlisides, & Mashour, 2022), and sleep (Brodbeck et al., 2012; Miyawaki, Billeh, & Diba, 2017; Simor et al., 2021). Microstate parameters were found to discriminate propofol-induced unconsciousness from regular wakefulness (Zhang et al., 2025), light and deep hypnosis (Katayama et al., 2007), and phases of transcendental meditation (Faber et al., 2017). Using EEG data recorded in 25 participants (Aicher et al., 2024), we tested whether neural metastability increases during a psychedelic state. We administered an ayahuasca-inspired formulation combining DMT and harmine to 25 participants while recording EEG data. We hypothesised that we might observe EEG microstate dynamics consistent with a more flexible and diverse sequence of brain states when the formulation is administered (relative to harmine and a placebo).

## Materials and methods

The methods and procedures for this study have been previously described in detail as part of a larger project (Aicher et al., 2024). The core methodology is summarized below.

### Participants

The final study sample consisted of 31 healthy male participants (mean age 25.4 ± 4.2 years) who completed all three intervention days. Participants were selected from an initial recruitment pool of 37 individuals. Six individuals withdrew prior to study completion. Inclusion criteria stipulated that only male participants could be included to avoid confounding effects of the menstrual cycle. Participants needed aged between 20 and 40 years, with a BMI of 18.5–30 kg/m². Volunteers were excluded if they had any current or past history of significant somatic, neurological, or psychiatric disorders, or a family history of severe psychiatric conditions such as psychosis. Further exclusion criteria included the use of medication that could interact with the study drug or any regular use of illicit substances (with a lifetime maximum of 15 psychedelic experiences). All participants agreed to abstain from alcohol the day before and caffeine on the day of each session, and to refrain from all psychoactive substance use for two weeks prior to and throughout the study period. Participants were monetarily compensated for their time. The study protocol was approved by the Cantonal Ethics Committee of Zürich (Basec-Nr. 2018–01385) and the Swiss Federal Office of Public Health. All procedures were conducted in accordance with the Declaration of Helsinki, and all participants provided written informed consent prior to inclusion.

### Experimental Design and Procedure

The study employed a randomized, double-blind, placebo-controlled crossover design. Each of the thirty-one participants completed three distinct experimental conditions on three separate intervention days, with a minimum two-week washout period between each session. The three conditions were: (1) 100 mg buccal harmine followed by 100 mg intranasal DMT (DMT/HAR); (2) 100 mg buccal harmine followed by intranasal placebo (HAR); and (3) buccal placebo followed by intranasal placebo (PLA). The order of these conditions was counterbalanced across participants.

On each intervention day, participants arrived at the laboratory in the late morning. Following their arrival, they completed baseline questionnaires before being fitted with measurement devices for continuous data collection, including high-density EEG, as well as an intravenous catheter for blood sampling. The administration of the study drug began at a fixed time (11:30 AM). The experimental procedure was identical for all three conditions. Data regarding the acute subjective experience were collected throughout the day, with retrospective questionnaires administered during the afterglow phase (approximately 300 minutes post-DMT administration). Online follow-up assessments were also conducted one and four months after the final intervention day.

### Setting and drug administration

All experimental sessions were conducted in soundproof, temperature-controlled laboratory rooms designed to create a comfortable, living-room-like atmosphere with dimmable lighting. No music was played during the experiment. Participants rested in a comfortable seated position throughout the experiment under the constant supervision of a medically and psychologically trained investigator.

For drug administration, this study utilized an ayahuasca-inspired formulation combining purified DMT and synthesized harmine. To enhance absorption kinetics and bypass first-pass metabolism, harmine was delivered via a 100 mg orodispersible tablet for buccal absorption, while DMT was administered via an intranasal spray to allow for incremental dosing. The dosing regimen began with the buccal administration of 100 mg harmine HCl or a matching placebo. Thirty minutes later, repeated intermittent intranasal dosing of DMT/HAR or placebo was initiated. The regimen consisted of ten 10 mg doses administered at 15-minute intervals over a 150-minute period, targeting a total cumulative dose of 100 mg of DMT/HAR. To ensure tolerability, participants had the option at each interval to skip a dose or reduce it to 5 mg. This option was exercised infrequently, resulting in only two participants receiving a slightly lower cumulative dose of 90 mg.

**Figure 1.**
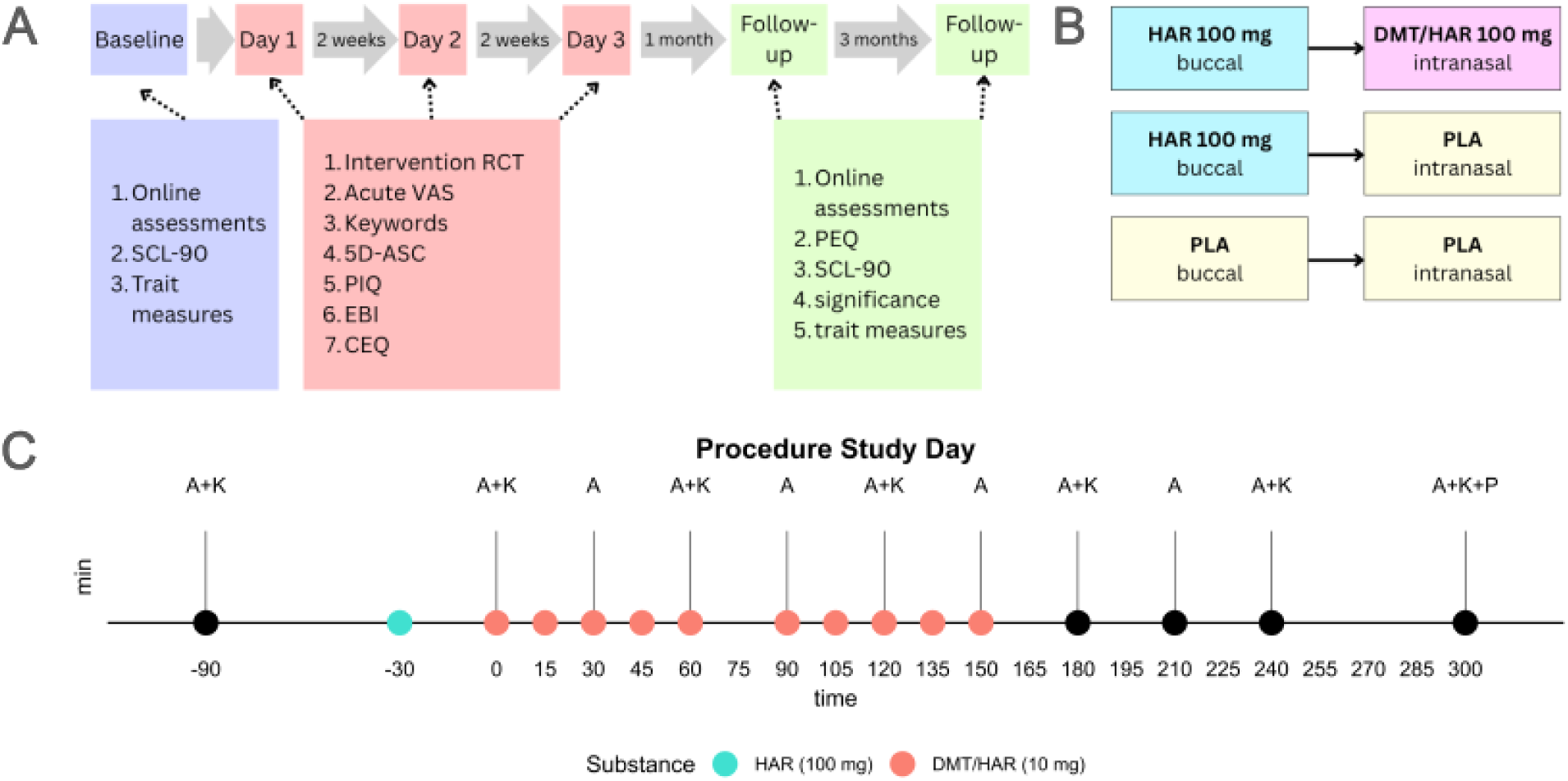
(A) Experiment Timeline. SCL-90-R: Symptom checklist; 5D-ASC = 5 dimensions altered states of consciousness rating scale; PIQ = psychological insights questionnaire; EBI = emotional breakthrough inventory; CEQ = challenging experience questionnaire; PEQ = persisting effects questionnaire; (B) Experimental conditions: DMT + HAR, HAR + PLA, PLA + PLA. (C) Timeline of a single testing day; A = acute assessment; K = keywords experience sampling; P = post-acute assessment (adapted from Aicher et al., 2024).

### Questionnaires

A comprehensive set of psychometric instruments was used to assess participants at multiple timepoints. The following measures, which are described in full detail in the original study publication (Aicher et al., 2024), were used to derive the outcomes for the present analysis. Questionnaires not available in German were adapted using a standard translation-back-translation methodology.

#### Baseline measures

Before the first intervention and at one- and four-month follow-ups, baseline psychological distress was assessed using the Symptom Check-List-90-Revised (SCL-90-R). Long-term changes attributed to the experiences were measured at follow-up with the Persisting Effects Questionnaire (PEQ), and the subjective significance of the experience was rated using questions adapted from Griffiths et al. (2006) (GSR). Several trait measures were also assessed at these timepoints, including well-being (WHO-5) and personality traits such as openness and neuroticism (IPIP).

#### Acute Experience Measures

The subjective effects during each session were tracked in real-time using a custom-developed experience sampling tool. At regular 30-minute intervals, participants used Visual Analog Scales (VAS) on a tablet to rate the intensity, valence (e.g., “liking,” “frightening”), and specific qualities of the experience, including items assessing elementary and complex visual imagery, and feelings of “letting go” versus “loss of control.”

Additionally, at 60-minute intervals, participants rated a set of 20 keywords developed to differentiate between “psychedelic” (e.g., “visuals,” “dreamlike”) and “empathogenic” (e.g., “open,” “connected”) phenomenological domains. The perceived integrity of body boundaries was also measured using the Perceived Body Boundaries Scale (PBBS).

#### Post-Acute Retrospective Experience Measures

Following the acute effects phase on each intervention day (at approx. t+300 min), the overall character of the altered state was retrospectively captured using the 11 subscales of the 5D-Altered States of Consciousness questionnaire (5D-ASC), which includes items such as “experience of unity,” “spiritual experience,” and “insightfulness.” Further retrospective measures, completed either on the same day or the following day, assessed the degree of acute psychological insight (Psychological Insights Questionnaire; PIQ), emotional release (Emotional Breakthrough Inventory; EBI), and the nature of any difficult aspects of the experience (Challenging Experience Questionnaire; CEQ). Qualitative diary reports were also collected to complement the quantitative data.

### Preprocessing

We collected EEG data using a 64-channel BioSemi active EEG system (BioSemi, Amsterdam, The Netherlands). EEG data were preprocessed using Matlab 2021b (The MathWorks, 2021) and EEGLAB (Delorme & Makeig, 2004). We initially filtered the data between 0.5 and 35 Hz and epoched the continuous data into segments with a length of one second. Then, we removed noisy trials and channels using a semi-automated pipeline in which trials and channels with a variance exceeding 90□V. were automatically rejected. Using Independent Component Analysis, we rejected EEG components likely representing movement or sweat artefacts. Having referenced the EEG data to the average, we reconstructed noisy channels using spherical spline interpolation. After cleaning, only 25 datasets were retained for further analysis. We only analysed data at t90 which corresponds to the time point at which the psychedelic experience was strongest.

### Microstate analysis

Microstate analysis was performed in Matlab 2021b using the +Microstate Toolbox (Tait & Zhang, 2022). In a first step, we used a k-means clustering algorithm to estimate sensor-space topographical maps. We identified local maxima in global field power (GFP) where we expect topographies to be stable. We sampled 3000 GFP peaks per participant and condition and stored the data in a cohort. K-means clustering assigns each data sample to a single microstate map using not more than 100 iteration and 20 replicates. We determined the optimal number of microstates k using the kneedle algorithm for 2-16 states. Finally, we fit the global output microstate maps back to the EEG data by assigning every data sample to the most similar global microstate map.

Microstates can be described using a range of microstate statistics. We calculated the average time the cortex spends in a microstate class (mean duration), the percentage of time the cortex stays in a microstate class (coverage), and the number of times a microstate class can be detected within a second (occurrence). By performing a spatial correlation between the global microstate maps and each EEG datapoint, we can obtain the Global Explained Variance (GEV) which is a metric describing how well each EEG data point is explained by the global microstate maps. GEV is calculated through an iterative process. For each microstate class, microstate maps from the time points in which spatial correlation with the current microstate class is highest are averaged and the new spatial correlation and GEV for the resulting microstate classes is obtained. This process is repeated until GEV is stable. Finally, we calculated the Hurst exponent which indexes long-range correlations in microstate sequences. For each microstate statistic, we performed a within-subject non-parametric Aligned Rank Transformed (ART) ANOVA to examine differences between DMT/HAR, HAR and PLA (with FDR corrections for multiple comparisons) based on previous work (Guo et al., 2025). Post-hoc testing was performed using ART contrasts. Analyses were performed using the Package ARTool version 0.11.2 in R version 4.4.2 (Kay, Elkin, Higgins, & Wobbrock, 2025) using RStudio Version 2025.05.0+496 (Posit team, 2025).

To test for differences in microstate topography between conditions, we performed a series of pairwise non-parametric permutation Topographical Analyses of Variance (TANOVA). TANOVAs were performed for each of the six microstate maps across the three condition pairs: DMT/HAR vs. PLA, DMT/HAR vs. HAR, and PLA vs. HAR. For each comparison, the global dissimilarity statistic (DISS) was calculated between the condition-averaged maps. Statistical significance was assessed against a null distribution generated from 5000 random permutations of the condition labels within each participant.

### Association between EEG microstates and subjective experience

To investigate how subjective psychedelic experience relates to changes in EEG microstate dynamics, we performed a mass-univariate analysis using Linear Mixed-Effects Models (LMMs) in R (v4.3.x) with the lme4 package. A separate LMM was fitted for each combination of microstate parameters and the 11 subscales of the ASD. In each model, the microstate parameter was predicted by the fixed effects of drug condition, the mean-centered questionnaire score, and their interaction. Drug condition was a factor with three levels (PLA/DMT/HAR), with PLA serving as the reference level. To account for non-independence of repeated measures from the same individual, a random intercept was included for each participant. Our primary analysis focused on whether the relationship between subjective experience and microstate dynamics was significantly different in the DMT/HAR condition compared to PLA. We applied a single False Discovery Rate (FDR) correction using the Benjamini-Hochberg procedure. An interaction was considered significant if its FDR-corrected p-value was less than 0.05.

## Results

### Microstate analysis

We used the kneedle algorithm to identify six global microstate classes best describing the temporal dynamics of the EEG dataset (Figure 2, A, C). First, we assessed global changes in microstate dynamics between the DMT/HAR, HAR and PLA conditions (Figure 2, B and Table 1 and 2). Evidenced by a shorter mean microstate duration (F(2, 48) = 32.04, p < .001), we found that the cortex transitions faster through microstate classes when DMT/HAR rather than HAR or PLA is administered. Post-hoc tests confirmed that the mean duration was shorter in the DMT/HAR condition compared to both HAR (t = −7.20, p < .001) and PLA (t = −6.63, p < .001). In line with this finding, we identified more local GPF maxima per second during the DMT/HAR experience relative to HAR (t = 6.18, p < .001) and PLA (t = 6.19, p < .001), with the main ANOVA showing a significant effect (F(2, 48) = 25.51, p < .001). DMT/HAR also produced more complex microstate topographies than HAR (t = 2.91, p = .017), with a significant main effect of condition on complexity (F(2, 48) = 4.65, p = .014).

**Figure 2.**
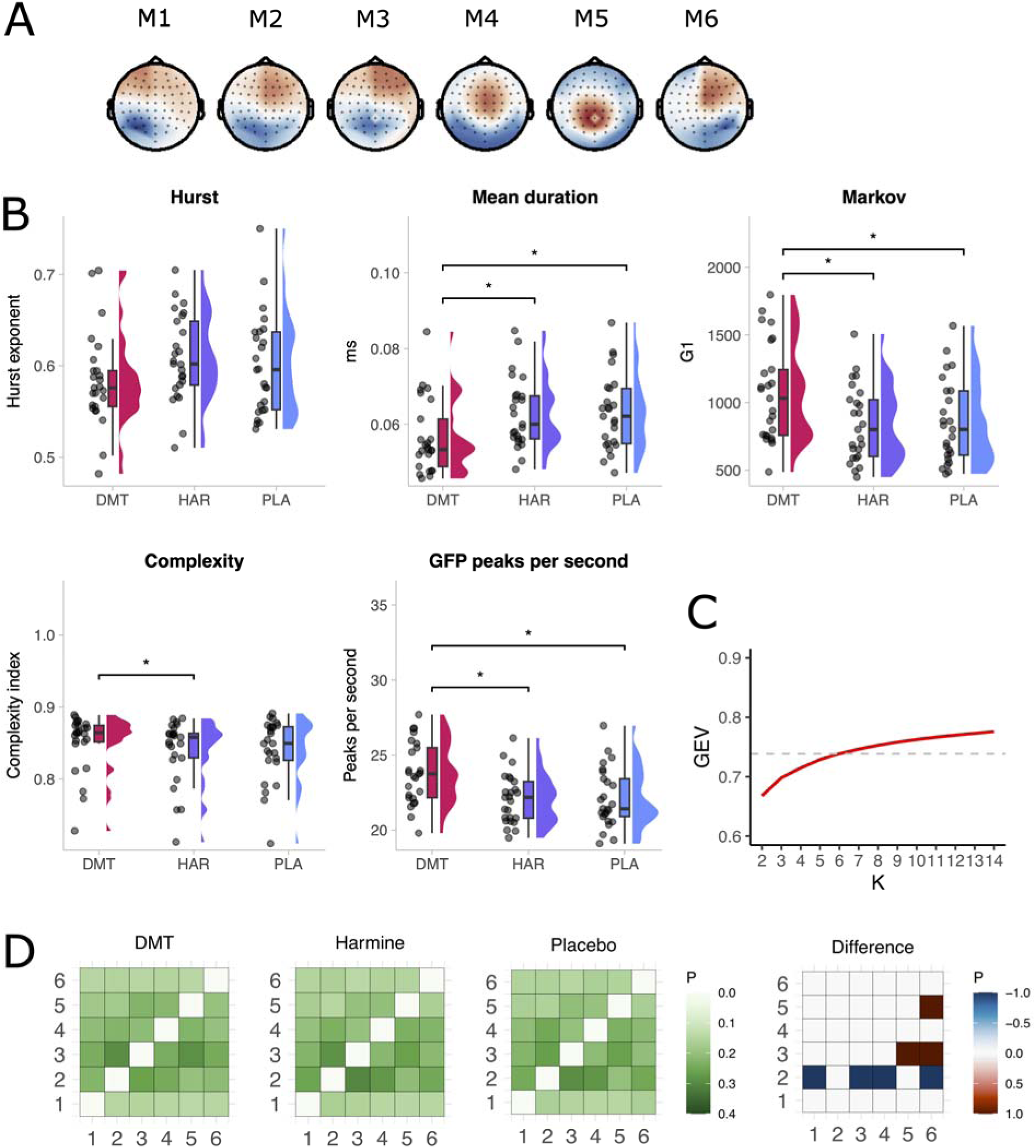
(A) Topographical maps of the six global microstate classes (M1-M6) identified by k-means clustering across all conditions. (B) Raincloud plots showing the distribution of global microstate parameters for DMT/HAR (red), HAR (purple), and PLA (blue) conditions. Asterisks indicate significant post-hoc comparisons (p < 0.05). (C) GEV as a function of the number of microstates K. A red dashed line flags K=6 as the optimal number of clusters identified with the kneedle algorithm. (D) Transition probability matrices for each condition and a difference matrix showing the change in transition probabilities between the DMT/HAR and PLA conditions. Warmer colors in the difference matrix indicate transitions that became more probable under DMT/HAR, while cooler colors indicate transitions that became less probable.

The first-order Markov statistic was higher in DMT/HAR than in the control conditions (main effect: F(2, 48) = 32.43, p < .001; post-hoc DMT/HAR vs HAR: t = 7.02, p < .001; post-hoc DMT/HAR vs PLA: t = 6.92, p < .001). To investigate long-range temporal correlations in the microstate sequences, we calculated the Hurst exponent. Across all three conditions, the Hurst exponent was consistently greater than 0.5, indicating the presence of persistent, long-range temporal correlations in the data. However, there was no main effect of condition on the Hurst exponent (F(2,48) = 2.85, p = .129), suggesting this long-range structure was not modulated by the drug administration. We performed a syntax matrix analysis to examine whether cortical topographies differ in their likelihood to transition from one microstate class to the next (Figure 2, D and Table 5 and 6).

Several key differences emerged in the DMT/HAR condition. The probability of transitioning to microstate M2 was significantly reduced from multiple other classes, including M1 (DMT/HAR vs PLA: t = −2.39, p = .042), M3 (DMT/HAR vs PLA: t = −2.39, p = .042), M4 (DMT/HAR vs HAR: t = −3.58, p = .002; DMT/HAR vs PLA: t = −3.46, p = .002), and M6 (DMT/HAR vs HAR: t = −4.39, p < .001; DMT/HAR vs PLA: t = −3.93, p < .001). In contrast, transitions towards microstates M3 and M5 were more likely. Specifically, during the DMT experience, cortical topographies are more likely to transition from microstate class M6 to M5 (DMT/HAR vs HAR: t = 3.37, p = .003; DMT/HAR vs PLA: t = 3.57, p = .002) and M3 (DMT/HAR vs HAR: t = 2.69, p = .020; DMT/HAR vs PLA: t = 3.43, p = .004), as well as from M5 to M3 (DMT/HAR vs HAR: t = 3.08, p = .010; DMT/HAR vs PLA: t = 2.57, p = .027).

Next, we analysed the specific parameters for each of the six microstates (Figure 3 and Table 5 and 6). The proportion of time covered by microstate M2 was significantly reduced under DMT compared to both HAR (t = −3.58, p = .002) and PLA (t = −2.65, p = .022). Conversely, coverage for microstates M3 and M5 was significantly increased during the DMT/HAR condition compared to controls (M3: DMT/HAR vs HAR t = 2.54, p = .029; DMT/HAR vs PLA t = 3.67, p = .002; M5: DMT/HAR vs HAR t = 4.04, p < .001; DMT/HAR vs PLA t = 3.98, p < .001). The average duration of all microstates except M3 was significantly shorter under DMT/HAR compared to both HAR and PLA (M1, M2, M4, M5, and M6; p < .01). Furthermore, the occurrence for microstates M3 and M5 was significantly higher in the DMT/HAR condition than in both control conditions (M3: DMT/HAR vs HAR t = 4.77, p < .001; DMT/HAR vs PLA t = 5.86, p < .001; M5: DMT/HAR vs HAR t = 6.37, p < .001; DMT/HAR vs PLA t = 6.04, p < .001).

**Figure 3.**
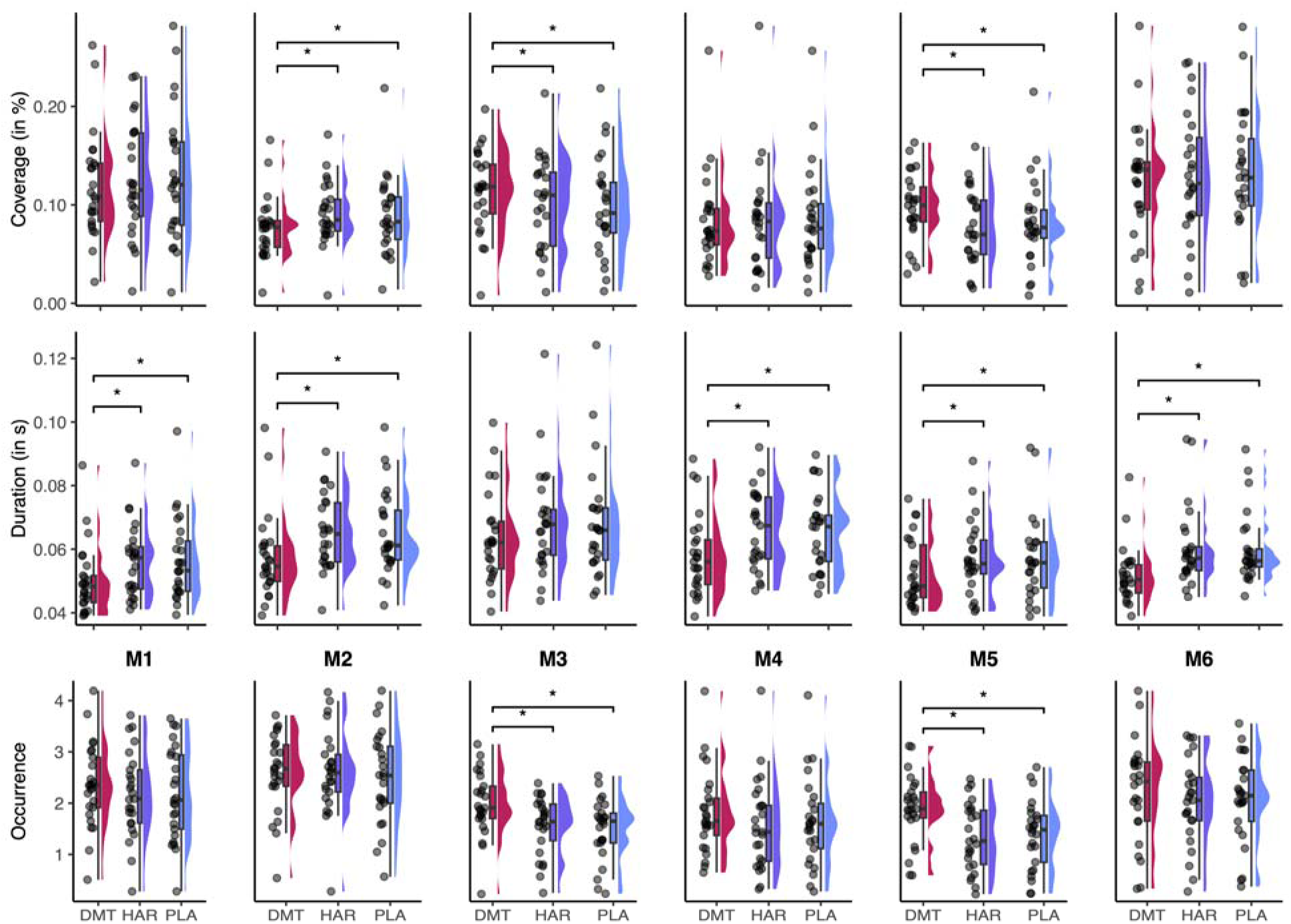
Raincloud plots showing the distribution of Coverage (top row), Duration (middle row), and Occurrence (bottom row) for each of the six microstates (M1-M6) across the DMT/HAR (red), HAR (purple), and PLA (blue) conditions. Asterisks indicate significant post-hoc comparisons (p<.05).

Using a TANOVA analysis, we revealed that the topographies of microstates M1, M2, M3, M4, and M6 were different in the DMT/HAR condition compared to both PLA and HAR (all p < .05). Microstate M5 was the only class that did not show a significant topographical change in any comparison. Importantly, no topographical differences were observed between PLA and HAR for any of the six microstates, confirming that the observed effects were specific to DMT/HAR administration (Figure 4 and Table 1). We did not identify questionnaire scores predicting microstate parameters (Supplementary table).

**Figure 4.**
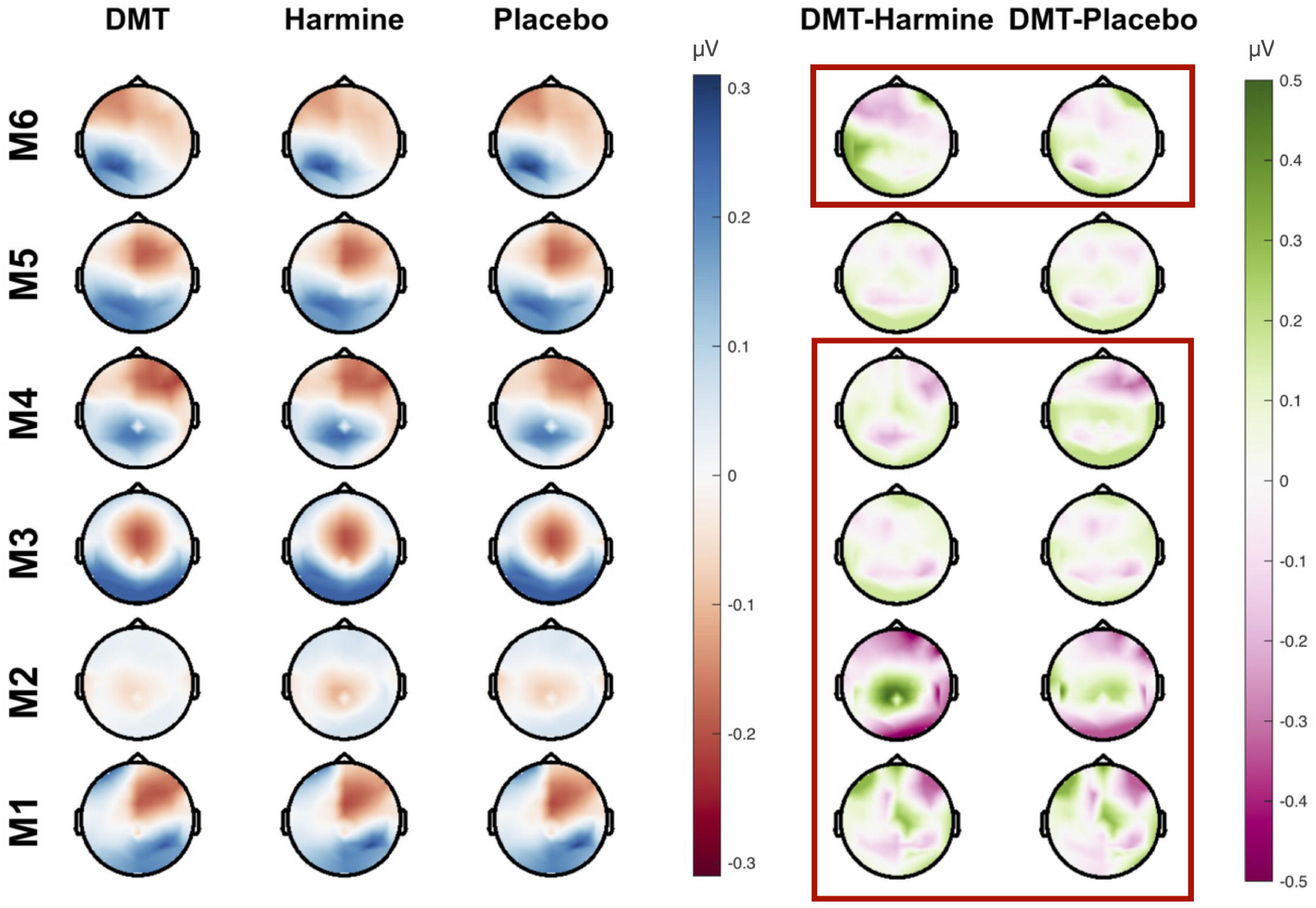
Topographical maps for each of the six microstates (M1-M6) are shown separately for the DMT/HAR, HAR, and PLA conditions. The final two columns show difference maps, illustrating the topographical changes induced by DMT relative to HAR and PLA. Red areas in the difference maps indicate regions with increased positive potential, while blue areas indicate regions with increased negative potential in the DMT/HAR condition. Significant differences between topographies are highlighted with a red rectangle.

## Discussion

Here we delineated the effects of DMT/HAR on the temporal dynamics of whole-brain functional states using EEG microstates analysis. Our results document a profound reconfiguration of microstate dynamics in psychedelic states induced by DMT/HAR. This was characterised by an acceleration of state transitions, an increase in topographical complexity, and a systematic restructuring of their underlying syntax. Our findings lend considerable weight to complex systems theories of psychedelic action, providing a detailed neurophysiological account of the DMT/HAR-induced state of consciousness.

Our findings refine dominant theories in psychedelic neuroscience which posit that psychedelics increase the diversity of neural states. Theoretical frameworks such as the Entropic Brain Hypothesis and the associated REBUS model propose that psychedelics destabilise predictive coding in the cortical hierarchy by relaxing the precision of high-level priors, thereby enabling a more unconstrained exploration of its functional repertoire (R. L. Carhart-Harris & Friston, 2019; R. Carhart-Harris et al., 2014; Singleton et al., 2022; Zeifman et al., 2025). Consistent with this view, we found that the topographic variance in our data was explained best by six microstates rather than the canonical four microstates found in previous work (Lehmann, Ozaki, & Pál, 1987). Our findings also demonstrate a marked acceleration of microstate dynamics, which was evidenced by a significant reduction in mean state duration and a corresponding increase in GFP peak frequency. Supporting the view that psychedelics facilitate neural state transitions (Singleton et al., 2022), our data also show that barriers to microstate transitions are lowered in the DMT/HAR-induced psychedelic state.

Crucially, our results provide direct evidence for the proposal that psychedelics induce a state of increased neural metastability (Girn et al., 2023). This suggests that psychedelics prevent the brain from settling into stable attractors, instead promoting the traversal of a greater volume of its state space per unit of time. This observation is consistent with fMRI findings of an expanded repertoire of functional connectivity patterns when other classic psychedelics are administered (Lord et al., 2019; Tagliazucchi et al., 2014). These dynamics lend strong support to the “flattened energy landscape” model of psychedelic action, wherein neural states become less stable and transitions between them become more frequent and fluid (Atasoy et al., 2017; Singleton et al., 2022). Although we observed an increased diversity of neural states under DMT/HAR, the temporal sequence of microstates was found to be less random. Indeed, microstate transitions became more syntactically structured and predictable, as evidenced by a higher first-order Markov statistic in the DMT/HAR condition. This indicates that the brain traverses its expanded state space with greater speed and fluidity, but this exploration is constrained by a more rigid underlying syntax. While the six microstates identified in our analysis cannot be directly mapped to the canonical four, the dynamic modulation of microstate sequences under DMT/HAR reveals a clear functional reorganization. We observed that microstate M2 was less prevalent under DMT/HAR, evidenced by reduced coverage in the DMT/HAR condition relative to HAR and PLA. Congruent with this finding, microstates M1, M3, M4, and M6 were less likely to transition to microstate M2 in the DMT condition. Conversely, we found an increased prevalence of microstates M3 and M5, which showed significantly greater coverage and occurrence in the DMT/HAR condition relative to HAR and PLA. This observation was complemented by a syntactic shift favouring transitions into these states. To clarify the neuroanatomical substrates of the observed microstate dynamics, combined EEG-fMRI investigations are necessary to map them onto core brain networks implicated in psychedelic action (e.g., limbic, salience, and default mode networks) (Kwan, Olson, Preller, & Roth, 2022).

Our finding of globally accelerated dynamics also differentiates the DMT/HAR state from other forms of spontaneous non-ordinary states of consciousness. For example, dream-like bizarre experiences in virtual reality were associated with altered microstate coverage, but no change in mean microstate duration was observed (Denzer, Diezig, Achermann, Mast, & Koenig, 2024). In a study investigating microstates during transcendental meditation, there were no or minor difference in microstate duration between meditation and rest (Faber et al., 2017). Unlike in the psychedelic state, another group found increased duration and coverage of a microstate linked to default mode network activity in meditators (Panda et al., 2016). Whereas in our dataset, DMT/HAR led to decreased microstate duration for most maps, deep hypnosis increased the duration of some microstates and decreased it for others (Katayama et al., 2007).

The present study has several limitations. We only included men as participants to avoid confounds arising from hormonal cycles, which might potentially reduce the generalisability of the dataset. While we documented a robust drug-induced shift in microstate dynamics, we did not find a link between questionnaire scores capturing psychedelic experience and changes in microstate parameters. We did not find direct correlations between microstate parameters and subjective experience, which raises several possibilities: the psychometric tools might not have captured the relevant aspects of phenomenology, the underlying relationship might be non-linear, or no such direct association exists. Finally, our microstate analysis focused only on data sampled during the peak of the psychedelic experience. As a result, we do not capture the dynamic evolution of brain activity during the onset, come-down, and afterglow phases of the experience. Further work needs to elucidate shifts in microstate parameters and their association with subjective changes throughout the entire course of the psychedelic experience.

In conclusion, we show that the DMT/HAR-induced psychedelic state is underpinned by a syntactic reconfiguration of EEG microstate sequences. Our results reconcile two seemingly opposing features of this state: (1) a faster, more fluid exploration of a diverse functional repertoire, characteristic of heightened metastability, and (2) a simultaneous increase in the predictability of state-to-state transitions. This finding moves beyond the entropic brain model by revealing that the psychedelic brain is not simply more random, but rather operates under a different and more structured temporal grammar. We propose that this syntactic restructuring might contribute to the neural mechanisms of psychedelic action.

## Notes

### Competing Interest Statement

The author(s) declared the following potential conflicts of interest with respect to the research, authorship, and/or publication of this article: MN, DS, and HAD declare that they have no conflict of interest. DAD and MS declare that they co-founded Reconnect Labs, an academic spin-off at the University of Zurich, focused on the development of psychedelic medicines for mental health. MJM declares he owns shares of Reconnect Labs.

### Funding

- Swiss National Science Foundation: P0ZHP1_191935
- Bioentrepreneur Fellowship UZH: BIOEF-18-009
- Forschungskredit PostDoc UZH: FK-18-052

